# The Brain Age Gap as a Predictor of Alcohol Initiation in Adolescence

**DOI:** 10.64898/2026.03.10.710953

**Authors:** H. Byrne, R. Visontay, E. K. Devine, N. E. Wade, J. Jacobus, A. J. Moore, L. M. Squeglia, L. Mewton

**Author notes:** **Corresponding author:** Hollie Byrne The Matilda Centre for Research in Mental Health and Substance Use, The University of Sydney, Sydney, NSW, Australia. Joint senior authors (L. M. Squeglia, L. Mewton).

## Abstract

**Background:** Growing evidence suggests regional and network-level brain imaging features in late childhood are predictive of alcohol use in adolescence. However, the directionality of these effects (i.e. whether they reflect accelerated or delayed neuromaturation) are mixed. We applied a Brain Age Gap Estimation (BrainAGE) model to examine whether overall deviations from typical brain aging trajectories are predictive of (1) alcohol initiation and (2) use behaviour (experimentation versus binge drinking) in adolescence.

**Methods:** Data from the Adolescent Brain Cognitive Development study release 6.0 were used. Baseline (ages 9-11) structural imaging features (cortical volume, area, and subcortical volume) were used to estimate BrainAGE. Alcohol use was determined using self-report data from the Substance Use Interview and Timeline Follow-Back across follow-ups (waves 1-6; ages 10-17). Logistic generalized mixed effects models examined whether BrainAGE predicted group status between (1) non-initiators (n=3,639) and initiators (n=1,176), and; (2) experimentation (at least one full drink, no binge episodes; n=461) and binge drinking (at least one episode; n=438).

**Results:** When adjusting for age, sex, and pubertal status, a one-standard-deviation decrease in BrainAGE (equivalent to 1.64 years) at baseline was associated with a 9.5% increase in odds of alcohol initiation in adolescence. However, this effect did not survive adjustment for sociodemographic and prior alcohol exposure covariates. Further, BrainAGE did not discriminate between experimentation and binge drinking.

**Conclusions:** Findings suggest BrainAGE in late childhood may reflect potential risk for alcohol initiation, but not behaviours, in adolescence. However, this association likely reflects complex interactions between brain structure and contextual factors, warranting further investigation.

## Introduction

Alcohol use in adolescence poses a significant public health concern. Recent data from the World Health Organization shows 57% of adolescents reporting alcohol use before age 16, and 20% reporting having experienced significant drunkenness (more than one binge drinking episode) by this age (1). Earlier alcohol initiation is associated with increased risk of adverse outcomes across the lifespan, including subsequent heavy drinking and substance use disorders (2,3), poorer academic performance (4), greater risk-taking behaviours (5), and increased mental health and social difficulties (6). Because adolescence is a sensitive period for neurodevelopment, alcohol exposure during this time can also disrupt maturation of neural systems involved in executive function, reward processing, and emotion regulation (7), making adolescent alcohol use particularly consequential. As a result, clarifying the risk factors and mechanisms underlying alcohol initiation is critical for informing prevention efforts and early intervention during this age period.

While the impacts of alcohol use on the brain have been well documented, evidence increasingly suggests distinct neurobiological characteristics may predate alcohol use. Specifically, research suggests aberrant development of subcortical and frontal systems prospectively predicts early initiation and escalation of alcohol use in adolescence (8), aligning with hypotheses suggesting neurobiological correlates of addiction may reflect both causes and consequences (9,10). This is supported by recent studies conducted in large, population imaging datasets such as the Adolescent Brain Cognitive Development (ABCD) study that have identified structural morphometric properties at both the regional (10,11) and network (12) level in late childhood (ages 9-11) which are predictive of alcohol use behaviours in early adolescence (up to age 15). However, the directionality of these effects – specifically, whether these observations reflect ‘accelerated’ or ‘delayed’ neuromaturation – have yet to be determined. Further, it remains unclear the extent to which alcohol use behaviours, particularly experimentation compared to binge drinking, are differentially related to structural brain properties.

Currently, few analytical methods are available to robustly assess maturation in the developing adolescent brain. One approach is brain age gap estimation (often abbreviated as ‘BrainAGE’ or ‘BAG’), a neuroimaging-based metric that computes the difference between an individual’s predicted brain age – derived from machine learning models applied to brain-imaging measures – and their chronological age. Notably, while these models do not provide an accurate prediction of maturation, BrainAGE can offer a general indicative measure of deviations from typical brain aging trajectories (13). In this context, a positive gap is interpreted as the brain appearing “older” than expected in comparison to the normative or control sample, while a negative gap points to the brain appearing “younger”. BrainAGE has been frequently applied within the adolescent and adult alcohol literature to investigate the impacts of alcohol use on deviations from normative developmental trajectories, which have typically identified a positive (higher) BrainAGE following chronic and/or heavy alcohol use (14). However, the predictive utility of BrainAGE in the initiation and type of alcohol use behaviour (experimentation vs. binge drinking) remains unknown.

Using data from the ABCD study, this work aims to investigate whether BrainAGE in late childhood (baseline; ages 9-11) can predict alcohol initiation in mid adolescence (wave 6; ages 15-17). Next, this work aims to examine whether baseline BrainAGE can differentiate between adolescents who have experimented with alcohol (defined here as consumption of a full drink but no binge drinking) compared to those who report at least one binge drinking episode.

## Methods and Materials

### Ethics approval and consent to participate

Data from the ABCD Study release 6.0 were used, which includes behavioral and biological data from 11,868 participants aged 9-11 years at baseline. Participants were recruited from 21 sites across the United States between September 2016 and November 2018, with follow-up visits conducted at yearly intervals. Parents/caregivers provided signed informed consent and all participants gave assent. The ABCD protocol was approved by the centralized institutional review board (IRB) at the University of California, San Diego and by the IRBs at the 21 sites. For secondary analysis of de-identified ABCD data in the current study, no additional ethical approval was required. All methods were performed in accordance with the relevant guidelines and regulations.

### Participants

Release 6.0 contains data from baseline through to wave 6 follow-up, with varying numbers of missed visits across follow-ups and data collection still ongoing at this stage. As data at the final available follow-up (wave 6; ages 15-17) aligns with the critical window for adolescent alcohol initiation (15), we included all available data through to wave 6 (n = 5,056).

Participants who had not initiated alcohol use were retained only if they had data at wave 6, whereas initiators were included if initiation occurred at any point prior to or at wave 6. To evaluate whether non-initiators had sufficient time to initiate, we compared their age distribution at wave 6 with the age-at-first-use distribution among initiators (*Figure S1*). The overlap in these distributions suggests that the non-drinkers included in these analyses were comparable in age to the alcohol initiation group, supporting the use of this approach for classification.

For BrainAGE calculation, participants were excluded if they were missing structural MRI data at baseline (n = 72), did not meet neuroimaging quality-control criteria (n = 472), or lacked complete covariate information required for harmonization of imaging data across scanners (n = 841; see *Image Processing*). For group classification, participants were excluded if they reported consuming a full alcoholic drink at baseline (n = 21); if non-drinking status within the non-initiator group could not be confirmed at wave 6 (i.e. missing data/not yet tested at wave 6; n = 5,647; see *Alcohol use classification*); if binge drinking status within the experimenter subgroup could not be determined at wave 6 (n = 277; see *Alcohol use classification*). Of the 11,868 participants recruited at baseline, 4,815 met inclusion criteria (*Figure 1*; n = 1,176 initiators).

**Figure 1.**
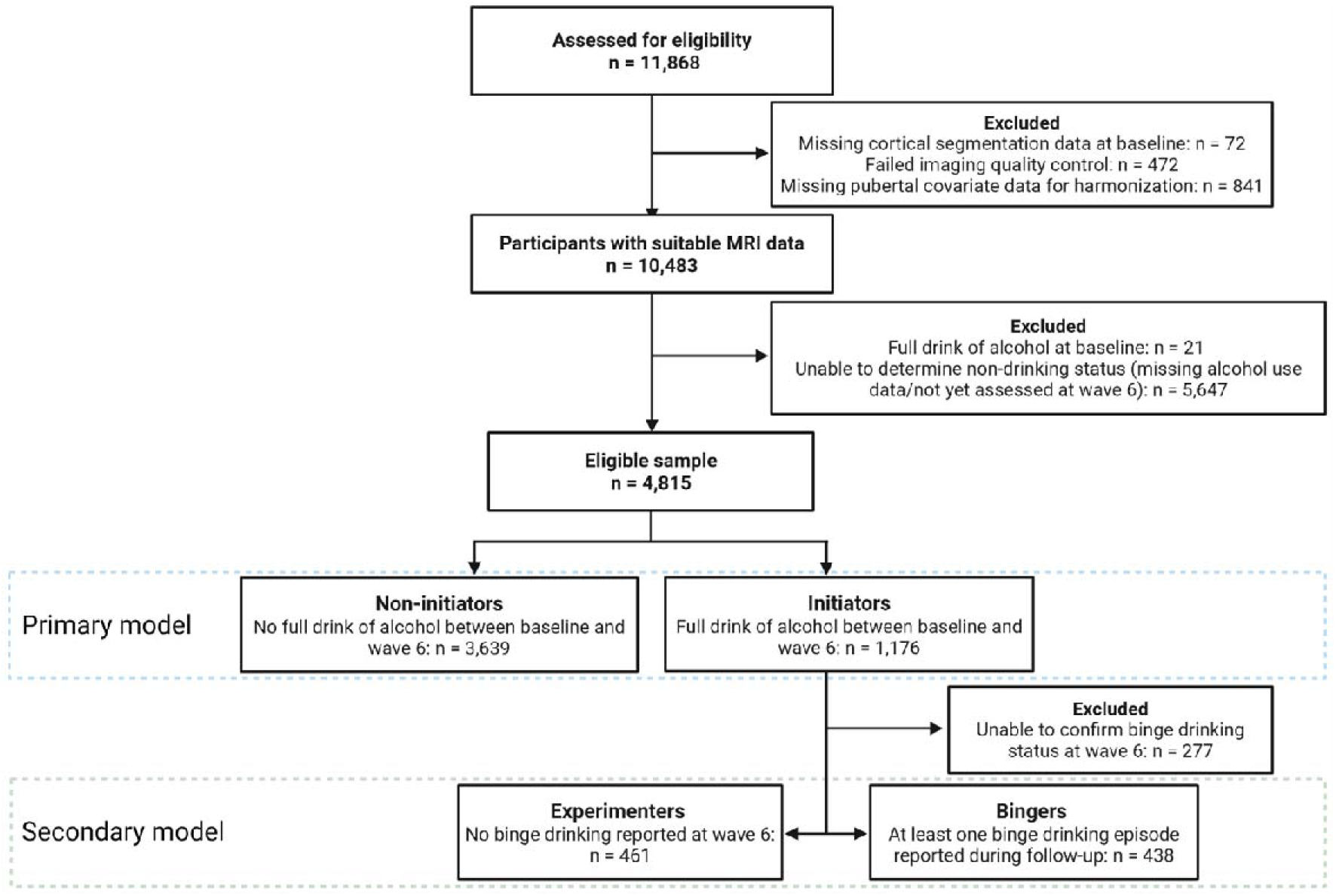
Flowchart for participant inclusion/exclusion criteria.

### Measures

#### Image processing

Preprocessed imaging data acquired at baseline (ages 9-11) were used (16). Cortical parcellation and subcortical segmentation of T1-weighted structural imaging data were performed using FreeSurfer (v7.1.1; https://surfer.nmr.mgh.harvard.edu/). Postprocessed cortical volume and area measurements mapped to the Desikan-Killiany atlas included 34 cortical parcellations per hemisphere, for a total of 68 regions of interest (ROIs) (17), respectively, and 53 bilateral global and subcortical volumetric measures derived from the FreeSurfer output.

To remove unwanted technical variability while preserving biological signals, neuroimaging measures were harmonized across scanners using ComBat (v1.0.13; Fortin et al., 2018). Scanner characteristics were modeled as batch covariates, while age, sex, and pubertal status (derived from parent-report Pubertal Development Scale scores; see *Pubertal status*) were preserved as biological covariates of interest.

#### BrainAGE model

To estimate BrainAGE, we used the Drobinin et al. (2022) model, which has demonstrated good external validity for use in adolescent datasets (20). The Drobinin model was trained on 1,299 participants aged 9–19 years, drawn from six independent non-ABCD samples. Age and sex distributions for each sample are reported in Drobinin et al. (2022). Participants were included in model training if they fell within the 9–19-year age range, passed automated MRI quality-control procedures, had no psychiatric diagnoses, and had an IQ greater than 75.

In Drobinin et al. (2022), model training was conducted within the *tidymodels* (version 0.1.1) framework, using the eXtreme Gradient Boosting (XGBoost) machine learning algorithm (version 1.0.0.2). Scan age in the Drobinin model was predicted from 189 features, including cortical gray matter volume and surface area measurements, as well as bilateral global and subcortical volume measurements. Cortical features were obtained using the FreeSurfer default Desikan-Killiany atlas, while bilateral global and subcortical volume measures were obtained using FreeSurfer segmentation output (aseg). The Drobinin model reported a mean absolute error (MAE; the average absolute difference between predicted and chronological age, where values of 1-2 reflect good model prediction accuracy in this context (13) of 1.53 years in typically developing adolescents; generalized to the validation dataset (mean absolute error = 1.55 years, 1.98 bias adjusted); and to the independent at-risk sample (mean absolute error = 1.49 years, 1.86 bias adjusted).

While the Drobinin et al. (2022) model was originally trained on 189 structural features, only 175 of these features were available in the ABCD dataset. Therefore, for the current study, the model was used to predict BrainAGE based on the 175 available features, replicating the methods by Whitmore et al. (2023). BrainAGE was estimated on the full harmonized sample (n = 10,483) prior to group classification.

#### Bias correction

Consistent with prior work (Le et al., 2018; Smith et al., 2019; Whitmore et al., 2023), we applied bias correction to account for systematic over- or underestimation related to age. Specifically, a linear model was fit to the validation data to derive a slope and intercept, which were then used to adjust predicted ages in the test set. Age was also included as a covariate in all subsequent statistical analyses to further control for residual age-related bias, as recommended in other work (13,24), and age outliers (participants aged ≤8.9 or ≥11.1; n = 3) were removed.

### Alcohol use classification

#### Alcohol use

Participants were classified into alcohol use groups using a hierarchical approach based on longitudinal self-reported data from the Substance Use Interview (SUI) and Timeline Follow-Back (TLFB) modules. Participants lacking complete SUI or TLFB data as per our exclusion criteria were excluded.

#### Primary model classification (non-initiators vs. initiators)

‘Non-initiators’ (n = 3,639) were defined as participants who did not report consumption of a full alcoholic drink in any session prior to and including wave 6. Participants with insufficient data at wave 6 to confirm non-initiation status at this timepoint were excluded. ‘Initiators’ (n = 1,176) were defined as participants who reported consuming at least one full alcoholic drink in any follow-up session before wave 6. This was assessed using binary ‘Yes’/‘No’ participant responses to the question: “Have you used since we last saw you: A full drink of beer, wine or liquor (rum, vodka, gin, whiskey),” as self-reported during the SUI.

#### Secondary model classification (experimenters vs. binge drinkers)

Among participants classified as ‘Initiators’, ‘Bingers’ (n = 438) were defined as participants who reported at least one binge drinking episode at any timepoint during follow-up. Binge episodes were defined as ≥4 standard drinks for females or ≥5 for males in a single drinking session, as self-reported in the TLFB. ‘Experimenters’ (n = 461) were defined as participants who reported a full drink of alcohol by wave 6 with no reported binge drinking episodes at any session prior to wave 6. To ensure Experimenters were appropriately characterized, Experimenters were required to have data available at wave 6 confirming non-binge drinking status at this timepoint (n = 277 excluded).

#### Covariates

Previous research has identified robust associations between age, sex, and pubertal status, respectively, in BrainAGE estimates (21,25). Therefore, these variables were included as primary covariates in the study. Chronological age and sex at baseline were determined from parent-reported date of child’s birth and sex assigned at birth, respectively. Baseline pubertal status was assessed using the parent-report Pubertal Development Scale (PDS), which evaluates physical markers of pubertal maturation such as growth of body hair, skin changes, and development of secondary sexual characteristics for males and females, respectively (26). An overall summary score ranging from 1 to 5 is calculated to represent each participant’s pubertal stage, approximating the Tanner stages of pubertal development ((27), with higher scores indicating more advanced pubertal development. Parent-report scores were used in this study based on recommendations for use of the PDS in baseline ABCD data (28).

### Statistical analysis

#### Primary analysis

A logistic generalized linear mixed effects model (GLMM) with a binomial distribution and logit link was used to predict alcohol use status (non-initiators vs. initiators) from baseline BrainAGE, controlling for baseline age, sex, and pubertal status (fixed effects), and random intercepts accounting for family nested within data collection site. Numeric predictors (BrainAGE, age, pubertal status) were z-standardized (mean = 0, SD = 1) for interpretability and effect size comparability. Models were fitted via maximum likelihood with Laplace approximation using the *lme4* (v 1.1.37; Bates et al., 2015) package in R (*bobyqa* optimizer). Effects are reported as odds ratios (ORs) with 95% confidence intervals. Significance was assessed with two-sided tests at α = .05.

#### Secondary analysis

The primary model was repeated using the same parameters to predict alcohol behaviour subgroup classification (experimentation vs. binge drinking) from baseline BrainAGE.

#### Sensitivity analyses

Two sensitivity analyses were performed using the primary model classification (non-initiators vs. initiators).

#### Sociodemographics and prior alcohol exposure

To assess whether observed differences in BrainAGE were independent of sociodemographic differences and/or prior alcohol exposure, the primary analysis was repeated with adjustment for covariates associated with brain structure and/or alcohol use in the ABCD sample. This included key sociodemographic variables (race, ethnicity, highest parental education, religious attitudes against alcohol), as well as additional variables related to prenatal exposure and adolescent alcohol use (maternal alcohol use during pregnancy, participant reported sipping at baseline), as performed in previous work (12). These variables were not interpreted as biological traits but included to reduce confounding related to structural and sociocultural factors known to affect neurodevelopmental trajectories and substance use risk. Further information on the variables used are reported in *Table S4*.

#### Exclusion of other substance use from non-initiator group

Inclusion of participants in the non-initiating group who initiate other substances during follow-up may confound results, as initiation of these substances may reflect shared high-risk neural substrates with alcohol (10). Therefore, to assess whether BrainAGE differs at baseline between ‘true’ non-initiators and those who later initiated other substances, the GLMM was repeated excluding non-initiators who initiated cigarette, e-cigarette, or cannabis smoking at baseline and/or during follow-up, as reported during the SUI (see *Table S4* for further detail on these measures).

## Results

### Participant characteristics

At baseline, participants who initiated alcohol use at or by wave 6 (n = 1,176) differed on nearly all characteristics compared to non-initiators (n = 3,639). Specifically, initiators were older than non-initiators, and a higher proportion were female, had been exposed to alcohol *in utero*, and reported alcohol sipping at baseline. In the non-initiating group, a greater proportion reported religious (negative) attitudes against alcohol compared to alcohol initiators, and groups differed in distribution of pubertal status, race, ethnicity, and highest caregiver education. Participant flowchart and characteristics are reported *Figure 1* and *Table 1*, respectively.

**Table 1.** Participant characteristics of full study sample.

| Characteristic | Non-initiators (n = 3,639) | Drinkers (n = 1,176) | p value |
| --- | --- | --- | --- |
| BrainAGE<sup>1,3</sup> | 0.52 (-0.56-1.70) | 0.30 (-0.76-1.52) | <0.001** |
| Age at baseline (years)<sup>1,3</sup> | 9.94 (9.41-10.48) | 10.35 (9.81-10.75) | <0.001** |
| Sex<sup>2,4</sup> |  |  | <0.001** |
| Female | 1,768 (49%) | 638 (54%) |  |
| Pubertal status at baseline<sup>2,4</sup> |  |  | <0.001** |
| 1 | 2,040 (56%) | 565 (48%) |  |
| 2 | 771 (21%) | 271 (23%) |  |
| 3 | 779 (21%) | 318 (27%) |  |
| 4 | 48 (1.3%) | 21 (1.8%) |  |
| 5 | <10 | <10 |  |
| Race<sup>2,5</sup> |  |  | <0.001** |
| American Indian/Alaska Native | 18 (0.5%) | <10 |  |
| Asian | 87 (2.4%) | 11 (0.9%) |  |
| Black (African American) | 454 (13%) | 55 (4.8%) |  |
| More than one race | 380 (10%) | 118 (10%) |  |
| Native Hawaiian/Other Pacific Islander | <10 | <10 |  |
| White | 2,610 (73%) | 944 (83%) |  |
| Missing/Other | 87 | 38 |  |
| Ethnicity (Hispanic)<sup>2,4</sup> | 663 (18%) | 215 (18%) | 0.9 |
| Missing | 46 | 10 |  |
| Caregiver education (highest)<sup>2,5</sup> |  |  | <0.001** |
| Up to high school (no diploma) | 131 (3.6%) | 25 (2.1%) |  |
| High school diploma/GED | 236 (6.5%) | 70 (6.0%) |  |
| Some college | 896 (25%) | 260 (22%) |  |
| Bachelor's degree | 1,045 (29%) | 288 (24%) |  |
| Graduate school or professional degree | 1,326 (36%) | 533 (45%) |  |
| Missing | <10 | 0 |  |
| Religious attitudes against alcohol<sup>2,4</sup> | 2,127 (59%) | 517 (44%) | <0.001** |
| Missing | 46 | 14 |  |
| Prenatal alcohol exposure<sup>2,4</sup> | 840 (24%) | 427 (39%) | <0.001** |
| Missing | 203 | 61 |  |
| Alcohol sipping at baseline<sup>2,4</sup> | 741 (20%) | 517 (44%) | <0.001** |
| Missing | 0 | 14 |  |

<sup>1</sup> Median (IQR); <sup>2</sup> n (%); <sup>3</sup> Wilcoxon rank sum test; <sup>4</sup> Pearson's Chi-squared test; <sup>5</sup> Fisher's exact test. Counts less than 10 have been suppressed to protect participant confidentiality. \* < 0.05; \*\* <0.001.

To assess whether baseline age differences could bias follow-up comparisons, where alcohol use might be skewed in the initiator group because of older age at baseline, age distributions for non-initiators at wave 6 and age at first reported drink in the initiator group were examined (*Figure S1*). No meaningful skew toward older ages was observed in the initiator group relative to non-initiators, suggesting that older age in the initiator group was unlikely to bias the overall findings.

In contrast to the full sample, when comparing drinking subgroup characteristics, Experimenters (n = 461) and Bingers (n = 438) only differed on a small number of measures (*Table 2*).

**Table 2.** Participant characteristics within experimenter and binge drinking subgroups.

| Characteristic | Experimenters (n = 461) | Bingers (n = 438) | p value |
| --- | --- | --- | --- |
| BrainAGE<sup>1,3</sup> | 0.32 (-0.78-1.55) | 0.19 (-0.83-1.41) | 0.5 |
| Age at baseline (years)<sup>1,3</sup> | 10.29 (9.74-10.73) | 10.42 (9.94-10.80) | 0.009* |
| Sex<sup>2,4</sup> |  |  | 0.2 |
| Female | 247 (54%) | 252 (58%) |  |
| Pubertal status at baseline<sup>2,4</sup> |  |  | 0.5 |
| 1 | 218 (47%) | 203 (46%) |  |
| 2 | 102 (22%) | 100 (23%) |  |
| 3 | 127 (28%) | 127 (29%) |  |
| 4 | 14 (3.0%) | <10 |  |
| 5 | 0 | <10 |  |
| Race<sup>2,5</sup> |  |  | 0.4 |
| American Indian/Alaska Native | <10 | <10 |  |
| Asian | <10 | <10 |  |
| Black (African American) | 23 (5.1%) | 16 (3.8%) |  |
| More than one race | 50 (11%) | 40 (9.5%) |  |
| Native Hawaiian/Other Pacific Islander | <10 | <10 |  |
| White | 373 (82%) | 283 (86%) |  |
| Missing/Other | <10 | 16 |  |
| Ethnicity (Hispanic)<sup>2,4</sup> | 63 (14%) | 87 (20%) | 0.013* |
| Missing | <10 | <10 |  |
| Caregiver education (highest)<sup>2,5</sup> |  |  | 0.068 |
| Up to high school (no diploma) | <10 | 14 (3.2%) |  |
| High school diploma/GED | 23 (5.0%) | 25 (5.7%) |  |
| Some college | 90 (20%) | 109 (25%) |  |
| Bachelor's degree | 108 (23%) | 110 (25%) |  |
| Graduate school or professional degree | 231 (50%) | 180 (41%) |  |
| Missing | 0 | 0 |  |
| Religious attitudes against alcohol<sup>2,4</sup> | 237 (52%) | 195 (45%) | 0.039* |
| Missing | <10 | <10 |  |
| Prenatal alcohol exposure<sup>2,4</sup> | 169 (40%) | 164 (40%) | 0.9 |
| Missing | 38 | 27 |  |
| Alcohol sipping at baseline<sup>2,4</sup> | 185 (40%) | 204 (47%) | 0.051 |
| Missing | 0 | 0 |  |

<sup>1</sup> Median (IQR); <sup>2</sup> n (%); <sup>3</sup> Wilcoxon rank sum test; <sup>4</sup> Pearson's Chi-squared test; <sup>5</sup> Fisher's exact test. Counts less than 10 have been suppressed to protect participant confidentiality. \* < 0.05; \*\* <0.001.

### BrainAGE model performance

In the original BrainAGE model validation, the model performed with a corrected (bias adjusted) MAE of 1.98 (Drobinin et al., 2022), aligning with the 1–2-year range typically reported by other adolescent BrainAGE models (30). In the current study, the model achieved an MAE of 2.28 prior to correction for age bias, and an MAE of 1.43 years after linear bias correction. These findings correspond closely to the study by Whitmore et al. using the same model in ABCD data (2.32 uncorrected, 1.45 uncorrected), with minor differences likely attributable to differences in harmonization following exclusion of participants with missing pubertal data. Similar to the findings by Whitmore et al. (2023), participants from the ABCD study also exhibited, on average, a marginally higher (more positive) BrainAGE compared to those included in the original Drobinin et al. (2022) model (*Table 2*).

### Baseline BrainAGE as a predictor of alcohol initiation

Baseline BrainAGE (*Figure 2*) significantly predicted alcohol use status by/at wave 6 (*Table 3*). Specifically, a one-standard-deviation increase (equivalent to 1.64 years) in BrainAGE was associated with an 8.7% decrease in the odds of alcohol use, holding other covariates constant. This equates to a 9.5% increase in the odds of alcohol use with a one-standard-deviation decrease in BrainAGE.

**Figure 2.**
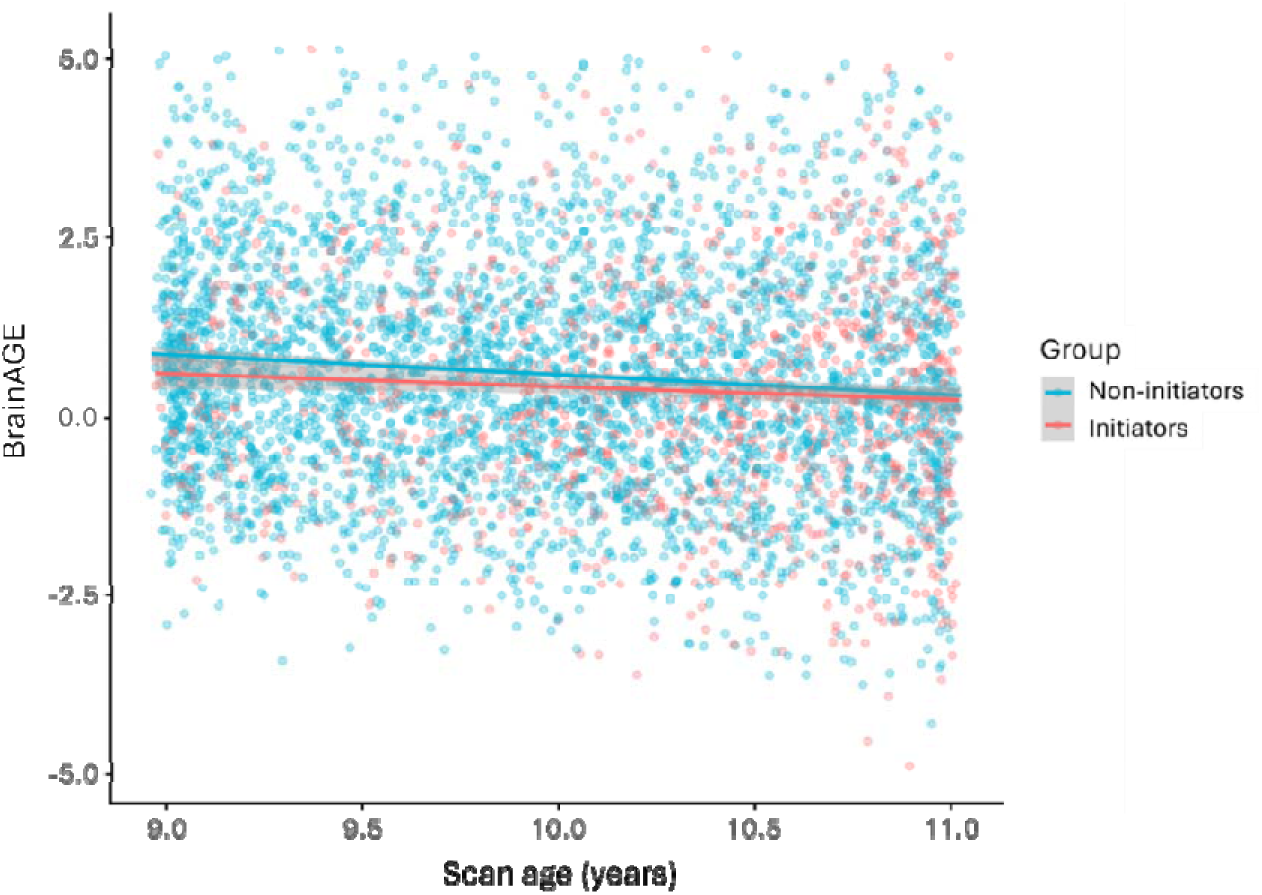
Scan age compared to BrainAGE within the full study sample.

**Table 3.** Results from logistic GLMM predicting alcohol use status (non-initiation vs. initiation) from baseline BrainAGE.

| Predictor | OR | 95% CI | *p* value |
| --- | --- | --- | --- |
| BrainAGE (std) | 0.913 | 0.844-0.987 | 0.023* |
| Age (std) | 1.7 | 1.55-1.86 | <.001** |
| Sex (female) | 1.28 | 1.06-1.54 | 0.011* |
| Pubertal status (std) | 1.04 | 0.95-1.14 | 0.377 |

OR = odds ratio; 95% CI = confidence intervals (odds ratio); std = standardized values. \* < 0.05; \*\* <0.001.

### Baseline BrainAGE as a predictor of alcohol use behaviours

Contrary to the primary model, BrainAGE was not identified as a significant predictor of drinking behaviour (experimentation versus binge drinking; *Figure 3*; *Table 4*).

**Figure 3.**
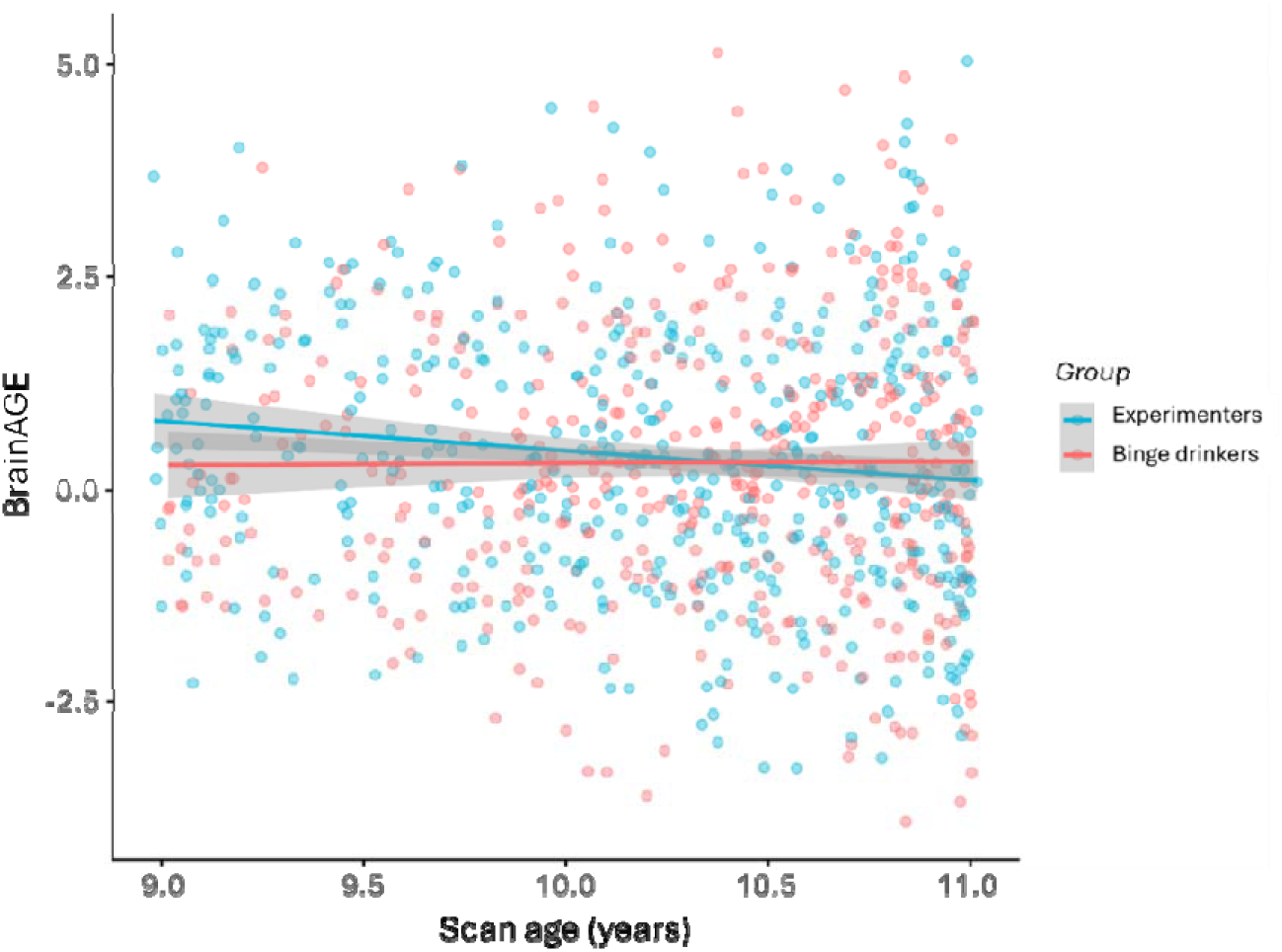
Scan age compared to BrainAGE between the experimenter and binge drinking groups.

**Table 4.** Results from logistic GLMM predicting alcohol use behaviour (experimentation vs. binge drinking) from baseline BrainAGE.

| Predictor | OR | 95% CI | *p* value |
| --- | --- | --- | --- |
| BrainAGE (std) | 0.999 | 0.862-1.158 | 0.989 |
| Age (std) | 1.235 | 1.048-1.455 | 0.012* |
| Sex (female) | 1.498 | 1.042-2.155 | 0.029* |
| Pubertal status (std) | 0.82 | 0.671-1.003 | 0.053 |

OR = odds ratio; 95% CI = confidence intervals (odds ratio); std = standardized values. \* < 0.05.

### Sensitivity analyses

#### Adjustment for secondary covariates

Following inclusion of secondary covariates in the primary model, no significant association between baseline BrainAGE and alcohol initiation status was identified (*Table S2*).

### Exclusion of participants of initiating other substances

Following exclusion of participants in the non-initiating group who initiated other substances at baseline and reported use at any time during follow-up, BrainAGE was still identified as a significant predictor of alcohol use (*Table S4*).

## Discussion

The present study investigated whether BrainAGE in late childhood could predict subsequent alcohol initiation and use behaviour in adolescence. Overall, ABCD participants showed a higher BrainAGE on average (i.e. more “mature appearing” brains) relative to chronological age compared to the model’s reference population. However, lower BrainAGE (a one-standard-deviation decrease) compared to the non-initiating group was associated with a 9.5% increased likelihood of alcohol initiation at follow-up, independent of chronological age, sex, and pubertal status. However, these effects were not robust to adjustment for sociodemographic characteristics and prior alcohol exposure. Additionally, within the initiation group, BrainAGE did not differentiate between experimentation and more harmful binge drinking behaviours. Overall, these findings suggest lower BrainAGE may predate alcohol initiation in adolescence, however, this relationship likely reflects complex, context-dependent risk rather than a direct effect of brain structure alone, warranting further investigation.

Results from our primary analysis suggest that lower BrainAGE (“less mature appearing” brains) in late childhood relative to non-initiators is predictive of subsequent alcohol initiation in adolescence. This stands in contrast to prior work identifying markers of accelerated neuromaturation, such as prefrontal cortical thinning (31,32) and regional volumetric increases (10), as predictors of alcohol use. This apparent discrepancy may reflect the limitations of region-of-interest approaches, given that brain development across childhood and adolescence is characterized by coordinated, network-level reorganization and structural co-maturation (33). By contrast, BrainAGE integrates cortical and subcortical features into a unified framework, where deviations reflect imbalances within and between these systems. This suggests that lower BrainAGE in the initiation group aligns with neurodevelopmental models proposing that such imbalances heighten vulnerability to substance use (34,35). However, collapsing these features into a summary score from a single timepoint precludes determination of how individual regions or systems contribute to the observed differences, and how these contributions evolve across development. Future work incorporating longitudinal and complementary network-level models alongside BrainAGE (such as structural covariance networks or connectome-based approaches) would help clarify whether disruption is coordinated or system-specific, more precisely mapping findings onto neurodevelopmental models of substance use vulnerability.

The attenuation of BrainAGE group differences following adjustment for sociodemographic characteristics and prior alcohol exposure suggests these factors may partially account for the observed relationship between BrainAGE and alcohol initiation, and warrants cautious interpretation of the primary findings. Both BrainAGE and alcohol initiation have been linked to a variety of environmental factors and life experiences, including socioeconomic status and childhood adversity (36–39), while the impacts of prenatal alcohol exposure on both regional and global brain development have been well-documented (40). This finding underscores the need to consider the contribution of biopsychosocial factors when examining neuroanatomical predictors of alcohol use, where future mediation and causal inference models will be essential for disentangling the independent and interactive contributions of these variables.

While BrainAGE was predictive of alcohol initiation, it did not differentiate between experimental and binge drinking behaviours, despite prior evidence that these may be neurobiologically distinct. For example, (41) demonstrated that structural brain features in early adolescence (age 14) are more strongly predictive of later binge drinking than experimental use, suggesting the neural correlates underlying escalation may be dissociable from and more pronounced than those associated with initiation alone. It is possible that a composite measure such as BrainAGE, when indexed at ages 9-11, may lack the sensitivity to capture such nuances, particularly given the reduced statistical power from a smaller binge drinking subsample. Additional factors including family history of substance use, impulsivity, and psychopathology have been linked to both binge drinking independently of experimentation (42,43) and to BrainAGE in adolescence (44,45), thus potentially obscuring group differences. Future work examining the relative contributions of such factors alongside BrainAGE would assist in determining its predictive utility across the spectrum of adolescent drinking behaviours.

## Limitations

Strengths of this work include the use of a large, demographically diverse prospective sample, consideration of multiple sociodemographic and alcohol exposure variables, and stringent alcohol use criteria. However, interpreting BrainAGE estimates in children and adolescents remains a significant challenge (Whitmore and Beck, 2025). As BrainAGE collapses hundreds of regionally distinct and asynchronously maturing features into a single global deviation score, such measures may obscure tissue- and network-specific patterns associated with genetic, environmental, and/or psychosocial factors. As these limitations are exacerbated within cross-sectional frameworks, normative BrainAGE trajectories in the paediatric population will be essential for informing interpretation of such metrics. While some longitudinal BrainAGE models are currently available (e.g. (21,36), the development of an externally validated, comprehensive model integrating multimodal imaging features will be pertinent to realizing this goal (20).

## Conclusion

The current findings suggest that BrainAGE in late childhood may reflect potential risk for subsequent alcohol use initiation, but not use behaviours, in adolescence. However, this association likely reflects complex, context-dependent risk rather than a direct effect of brain structure alone. Further research tracking longitudinal trajectories of BrainAGE within a latent change score framework to examine bidirectional relationships between other genetic, environmental, and psychosocial variables will be essential for determining the role of BrainAGE as a predictor of alcohol use in adolescence.

## Acknowledgements

Data used in the preparation of this article were obtained from the Adolescent Brain Cognitive Development^™^ (ABCD) Study, held in the NIH Brain Development Cohorts Data Sharing Platform. This is a multisite, longitudinal study designed to recruit more than 10,000 children age 9-10 and follow them over 10 years into early adulthood. The ABCD Study® is supported by the National Institutes of Health and additional federal partners under award numbers U01DA041048, U01DA050989, U01DA051016, U01DA041022, U01DA051018, U01DA051037, U01DA050987, U01DA041174, U01DA041106, U01DA041117, U01DA041028, U01DA041134, U01DA050988, U01DA051039, U01DA041156, U01DA041025, U01DA041120, U01DA051038, U01DA041148, U01DA041093, U01DA041089, U24DA041123, U24DA041147. A full list of supporters is available at https://abcdstudy.org/federal-partners.html. A listing of participating sites and a complete listing of the study investigators can be found at https://abcdstudy.org/consortium_members/. ABCD consortium investigators designed and implemented the study and/or provided data but did not necessarily participate in the analysis or writing of this report. This manuscript reflects the views of the authors and may not reflect the opinions or views of the NIH or ABCD consortium investigators. This work was supported by funding from the NIH (R01AA030575, Mewton/Squeglia; and K24AA031052, Squeglia).

## Author contributions

H.B. contributed to the conceptualization, data curation, formal analysis, visualization, and writing – original draft, reviewing and editing of the manuscript. L.M. and L.S. contributed to the conceptualization, supervision, interpretation of results, funding acquisition, and writing – reviewing and editing of the manuscript. R.V. and E.D. contributed to conceptualization, interpretation of results, and writing – reviewing and editing. N.E.W., J.J. and A.J.M. contributed to interpretation of results and writing – reviewing and editing. All authors have read and approved the final manuscript.

