## Supplementary tables and figures for "The Brain Age Gap as a Predictor of Alcohol Initiation in Adolescence"

**Supplementary Material**

*Age distributions*

**Figure S1.** Age distribution of non-initiators at wave 6 compared to age of reported initiation.

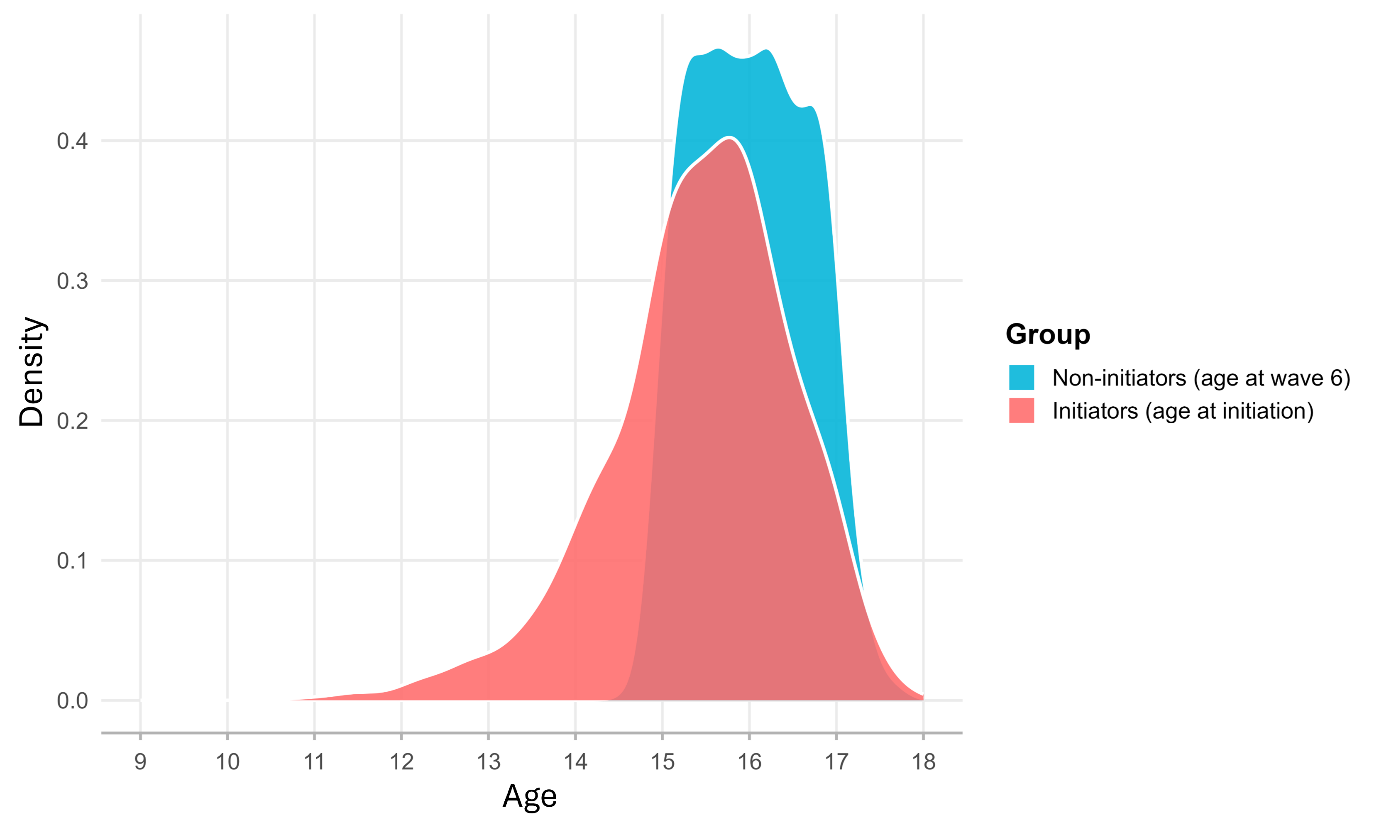

*Secondary covariate adjustment*

**Table S1.** Description and formatting of secondary covariates, each acquired from baseline.

| **Characteristic** | **Variable** | **Variable description** |
| --- | --- | --- |
| Race | ab_g_stc__cohort_race__nih | 7-level classification of race based on National Institutes of Health (NIH) standards: “American Indian/Alaska Native” / “Asian” / “Black (African American)” / “More than one race” / “Native Hawaiian/Other Pacific Islander” / “White” / “Other/Unknown”. |
| Ethnicity | ab_p_demo__ethn_001 | 4-level variable: “Do you consider the child Hispanic/Latino/Latina?”: “No” / “Yes” / “Decline to Answer” / “Don’t know”. |
| Caregiver education | ab_g_dyn__cohort_edu__cgs | 5-level cohort description of the highest education across caregivers: “Up to high school (No diploma)” / “High school diploma/GED” / “Some college” / “Bachelor’s degree” / “Graduate school or professional degree”. |
| Religious attitudes towards alcohol | ab_p_demo__relig__alc_001 /  ab_p_demo__relig_001 | 3-level variable: "Does your child's religion have rules forbidding the use of alcohol?" “No” / “Yes” / “Don’t know”. To account for missing data at baseline, these were populated with responses provided at next follow-up (wave 1), or coded as 0 (‘No’) if participants responded as not religious to “What is the child's religious preference?” (“Atheist” / “Agnostic” / “Nothing in particular”) at baseline. |
| Alcohol sipping at baseline | su_y_sui__use__alc__sip_001 /  su_y_sui__hrd__alc_001 | 2-level variable: "Have you ever used: A sip of alcohol such as beer, wine or liquor (rum, vodka, gin, whiskey)", “No” / “Yes”. NAs were populated with 0’s (“No”) in instances where participants responded “No” to “Have you heard of: alcohol, such as beer, wine or liquor?”. |
| Prenatal alcohol exposure | ph_p_dhx__alc_001a /  ph_p_dhx__alc_001b | Binary variable derived from caregiver responses to the following questions: “Before biological mother found out she was pregnant, but while she might have been pregnant with this child, did she use any of the following: Alcohol?” / “After biological mother found out she was pregnant, did she use any of the following: Alcohol?” (“No” / “Yes” / “Don’t know”). Participants who responded “yes” to either or both questions were coded as 1, participants who responded “no” to both questions were coded as 0. |

After introducing secondary covariates, models failed to converge. Diagnostic testing indicated that convergence issues were driven by the family ID random effect, which was affected by sparse clustering (i.e., many families containing only a single participant). To address this issue, siblings were excluded at random from the covariate analysis (n = 693) while maintaining site as a random effect.

**Table S2.** Results from logistic GLMM predicting early alcohol use status from baseline BrainAGE adjusting for baseline secondary covariates.

| **Predictor** | **OR** | **95% CI** | ***p* value** |
| --- | --- | --- | --- |
| BrainAGE (std) | 0.968 | 0.891-1.051 | 0.438 |
| Age (std) | 1.612 | 1.475-1.763 | <.001** |
| Sex (female) | 1.205 | 0.989-1.469 | 0.065 |
| Pubertal status | 1.135 | 1.025-1.257 | 0.015* |
| Race |  |  |  |
| Black/African American | 0.427 | 0.296-0.615 | <.001** |
| Asian | 0.388 | 0.185-0.814 | 0.012* |
| American Indian/Alaska Native | 1.817 | 0.63-5.235 | 0.269 |
| Native Hawaiian/Other Pacific Islander | 1.979 | 0.159-24.697 | 0.596 |
| More than one race | 0.946 | 0.721-1.24 | 0.686 |
| Ethnicity (non-Hispanic) | 0.789 | 0.613-1.016 | 0.066 |
| Caregiver education (highest) |  |  |  |
| High school diploma/GED | 2.002 | 1.006-3.985 | 0.048* |
| Some college | 1.774 | 0.951-3.31 | 0.072 |
| Bachelor’s degree | 1.389 | 0.74-2.608 | 0.307 |
| Graduate school or professional degree | 1.948 | 1.042-3.644 | 0.037 |
| Religious attitudes towards alcohol (yes) | 0.712 | 0.601-0.843 | <.001** |
| Alcohol sipping at baseline (yes) | 2.709 | 2.275-3.225 | <.001** |
| Prenatal alcohol exposure (yes) | 1.602 | 1.343-1.91 | <.001** |

OR = odds ratio; 95% CI = confidence intervals (odds ratio); std = standardized values.

*Other substance use*

After alcohol, cigarettes, e-cigarettes (vapes), and cannabis are the most commonly used substances among adolescents, with smoking or vaping as prevalent methods of consumption (Charrier et al., 2024). Therefore, participants who initiated these substances in the non-alcohol-initiating group were excluded from the analysis, using measures listed in *Table S4*.

**Table S3.** Description of variables used to define other substances in non-initiating group.

| **Characteristic** | **Variable** | **Description** |
| --- | --- | --- |
| Blunt/joint use | su_y_sui__use__mj__blunt_001 /  su_y_sui__use__mj__blunt_001__l | Binary Y/N response to the question: “Have you ever used/Since we last saw you, have you used: Blunts, when you combine tobacco and marijuana in joints.” |
| Cannabis use (smoking) | su_y_sui__use__mj__smoke_001 /  su_y_sui__use__mj__smoke_001__l | Binary Y/N response to the question: “Have you ever used/Since we last saw you, have you used: Smoked marijuana, also called pot, grass, weed, ganja - more than just a puff?” |
| Cigarette use | su_y_sui__use__nic__cig_001 /  su_y_sui__use__nic__cig_001__l | Binary Y/N response to the question: “Have you ever used/Since we last saw you, have you used: Tobacco cigarette?” |
| E-cigarette/vape use | su_y_sui__use__nic__vape_001 /  su_y_sui__use__nic__vape_001__l | Binary Y/N response to the question: “Have you ever used/Since we last saw you, have you used: Electronic nicotine or vaping products, such as an e-cigarette, vape pen, or Zen?” |

**Table S4.** Comparison of other substance use initiation between groups.

| **Characteristic** | **Non-initiators (n = 3,639)** | **Initiators (n = 1,176)** | ***p* value** |
| --- | --- | --- | --- |
| Substance use^1,2^ |  |  | <.001** |
| Joint/blunt | <10 | <10 |  |
| Cigarette | <10 | 12 (1.0%) |  |
| E-cigarette/vape | 186 (5.1%) | 162 (14%) |  |
| Cannabis | 58 (1.6%) | 83 (7.1%) |  |
| Multiple | 161 (4.4%) | 479 (41%) |  |

^1^ n (%); ^2^ Fisher’s Exact Test for Count Data with simulated p-value (based on 10000 replicates). ** <0.001.

**Table S5.** Results from logistic GLMM predicting alcohol use status from baseline BrainAGE after exclusion of other substances from the non-initiation group.

| **Predictor** | **OR** | **95% CI** | ***p* value** |
| --- | --- | --- | --- |
| BrainAGE (std) | 0.922 | 0.85-1.0 | 0.049* |
| Age (std) | 1.744 | 1.588-1.915 | <.001** |
| Sex (female) | 1.295 | 1.067-1.572 | 0.009* |
| Pubertal status (std) | 1.084 | 0.97-1.21 | 0.155 |

OR = odds ratio; 95% CI = confidence intervals (odds ratio); std = standardized values. * < 0.05; ** <0.001.
